# High throughput and affordable genome-wide methylation profiling of circulating cell-free DNA by Methylated DNA sequencing (MeD-seq) of LpnPI digested fragments

**DOI:** 10.1101/2021.07.12.452012

**Authors:** Teoman Deger, Ruben G Boers, Vanja de Weerd, Lindsay Angus, Marjolijn MJ van der Put, Joachim B Boers, Zakia Azmani, Wilfred FJ van Ijcken, Dirk J Grünhagen, Lisanne F van Dessel, Martijn PJK Lolkema, Cornelis Verhoef, Stefan Sleijfer, John WM Martens, Joost Gribnau, Saskia M Wilting

## Abstract

**Background:** DNA methylation detection in liquid biopsies provides a highly promising and much needed means for real-time monitoring of disease load in advanced cancer patient care. Compared to the often-used somatic mutations, tissue- and cancer-type specific epigenetic marks affect a larger part of the cancer genome and generally have a high penetrance throughout the tumour. Here we describe the successful application of the recently described MeD-seq assay for genome-wide DNA methylation profiling on cell-free DNA (cfDNA). The compatibility of the MeD-seq assay with different types of blood collection tubes, cfDNA input amounts, cfDNA isolation methods, and vacuum-concentration of samples was evaluated using plasma from both metastatic cancer patients and healthy blood donors (HBDs). To investigate the potential value of cfDNA methylation profiling for tumour load monitoring, we profiled paired samples from 8 patients with resectable colorectal liver metastases (CRLM) before and after surgery.

**Results:** The MeD-seq assay worked on plasma-derived cfDNA from both EDTA and CellSave blood collection tubes when at least 10 ng of cfDNA was used. From the 3 evaluated cfDNA isolation methods, both the manual QIAamp Circulating Nucleic Acid Kit (Qiagen) and the semi-automated Maxwell® RSC ccfDNA Plasma Kit (Promega) were compatible with MeD-seq analysis, whereas the QIAsymphony DSP Circulating DNA Kit (Qiagen) yielded significantly fewer reads when compared to the QIAamp kit (P<0.001). Vacuum-concentration of samples before MeD-seq analysis was possible with samples in AVE buffer (QIAamp) or water, but yielded inconsistent results for samples in EDTA-containing Maxwell buffer. Principal component analysis showed that pre-surgical samples from CRLM patients were very distinct from HBDs, whereas post-surgical samples were more similar. Several described methylation markers for colorectal cancer monitoring in liquid biopsies showed differential methylation between pre-surgical CRLM samples and HBDs in our data, supporting the validity of our approach. Results for MSC, ITGA4, GRIA4, and EYA4, were validated by quantitative methylation specific PCR.

**Conclusions:** The MeD-seq assay provides a promising new method for cfDNA methylation profiling. Potential future applications of the assay include marker discovery specifically for liquid biopsy analysis as well as direct use as a disease load monitoring tool in advanced cancer patients.

## BACKGROUND

Liquid biopsies, referring to the sampling of bodily fluids instead of tissue, provide a novel approach for real-time cancer screening, disease monitoring, and treatment selection in advanced cancer patients (reviewed in (1)). Currently, treatment of patients with advanced or metastatic solid tumours is more or less a trial-and-error process, guided where possible by molecular features of the (primary) tumour. Treatment response in patients is monitored by relatively insensitive and expensive imaging techniques, such as CT scans and MRI scans, on which (the lack of) treatment effect usually only becomes visible after 3 months or more. This leads to unnecessary toxicity, loss of valuable time when treatment is ineffective, and additionally causes anxiety for patients.

Liquid biopsies are known to contain trace amounts of cell-free DNA (cfDNA) and cells derived from the tumour and can be obtained on a regular basis in a minimally invasive manner (2). Interestingly, in patients with metastatic lung cancer the amount of circulating tumour DNA (ctDNA) in the blood was found to correlate with the total disease load, indicating that ctDNA can indeed be used to monitor disease progression (3).

However, for reliable estimation of tumour load, we need to ensure that the tumour-specific signal detected in the liquid biopsy is homogeneously present in virtually all cells of the tumour. This prohibits the use of polyclonal mutations, which greatly reduces the number of informative mutations for tumour load estimation in cfDNA (4); (5). Next to the heterogeneity within a tumour, heterogeneity between tumours further complicates the use of universal hotspot mutation panels for this purpose. In addition, clonal hematopoiesis, a common aging-associated phenomenon, can also generate mutations detectable in cfDNA, thereby obscuring the tumour-specific signal (6); (7). Finally, a large part of the tumour genome does not carry mutations and is therefore ignored when only mutations are investigated in cfDNA. Taking all this into consideration, other tumour-specific alterations like chromosomal copy number variations (CNVs) and DNA methylation changes may represent more promising markers for disease load estimation as they affect a much larger part of the tumour genome. Tumour-specific DNA methylation frequently occurs at CpG islands and is associated with silencing of tumour suppressor genes in cancer (reviewed in (8)). These alterations in DNA methylation are known to occur early on during cancer development and will therefore have a high penetrance throughout the tumour. In addition, different cell types have their own lineage-specific DNA methylation pattern, which is very different in epithelial cells compared to leukocytes, the predominant contributors to cfDNA (9). As a result, the signal-to-noise ratios for this type of molecular marker in blood-derived cfDNA will be even further enhanced. Indeed, results from the Circulating Cell-free Genome Atlas (CCGA; NCT02889978) study indicate that cfDNA methylation analyses outperform whole-genome sequencing (WGS) and targeted sequencing approaches interrogating CNVs and mutations for both the detection and classification of many cancer types (10).

The fact that most blood samples only yield small amounts of fragmented cfDNA combined with the low recovery of DNA after bisulfite conversion (22-66%) (11), renders conventional genome-wide DNA methylation analyses like whole genome bisulfite sequencing (WGBS) and reduced representation bisulfite sequencing (RRBS) on cfDNA technically challenging. However, a recent proof-of-principle study in paediatric solid tumours demonstrates the feasibility of an adapted RRBS protocol for cfDNA analyses (12); (13). Similarly, Shen et al. developed a method for high throughput sequencing of immune-precipitated methylated cfDNA (cfMeDIP-seq), which uses antibody-based enrichment of methylated DNA sequences and is therefore not dependent on bisulfite treatment (14). However, the use of antibodies invariably introduces noise. In addition, a bias for CpG-poor regions was observed for MeDIP approaches before, resulting in an overall coverage of about 16% of all CpG dinucleotides genome-wide (15).

Therefore, we applied the recently described MeD-seq method (16), which uses the LpnPI methylation-dependent restriction enzyme to allow for genome-wide methylation profiling without the need for harsh bisulfite treatment or the use of potentially biased enrichment techniques, to cfDNA samples (Figure 1). Contrary to other methylation-dependent enzymes, LpnPI activity is restricted by a short template size, which prevents over-digestion of CpG-dense regions and subsequent mapping problems. MeD-seq was shown to reliably detect DNA methylation at >50% of all CpG dinucleotides genome-wide without a preference for CpG-dense or CpG-poor regions at a low sequencing depth (16).

**Figure 1.**
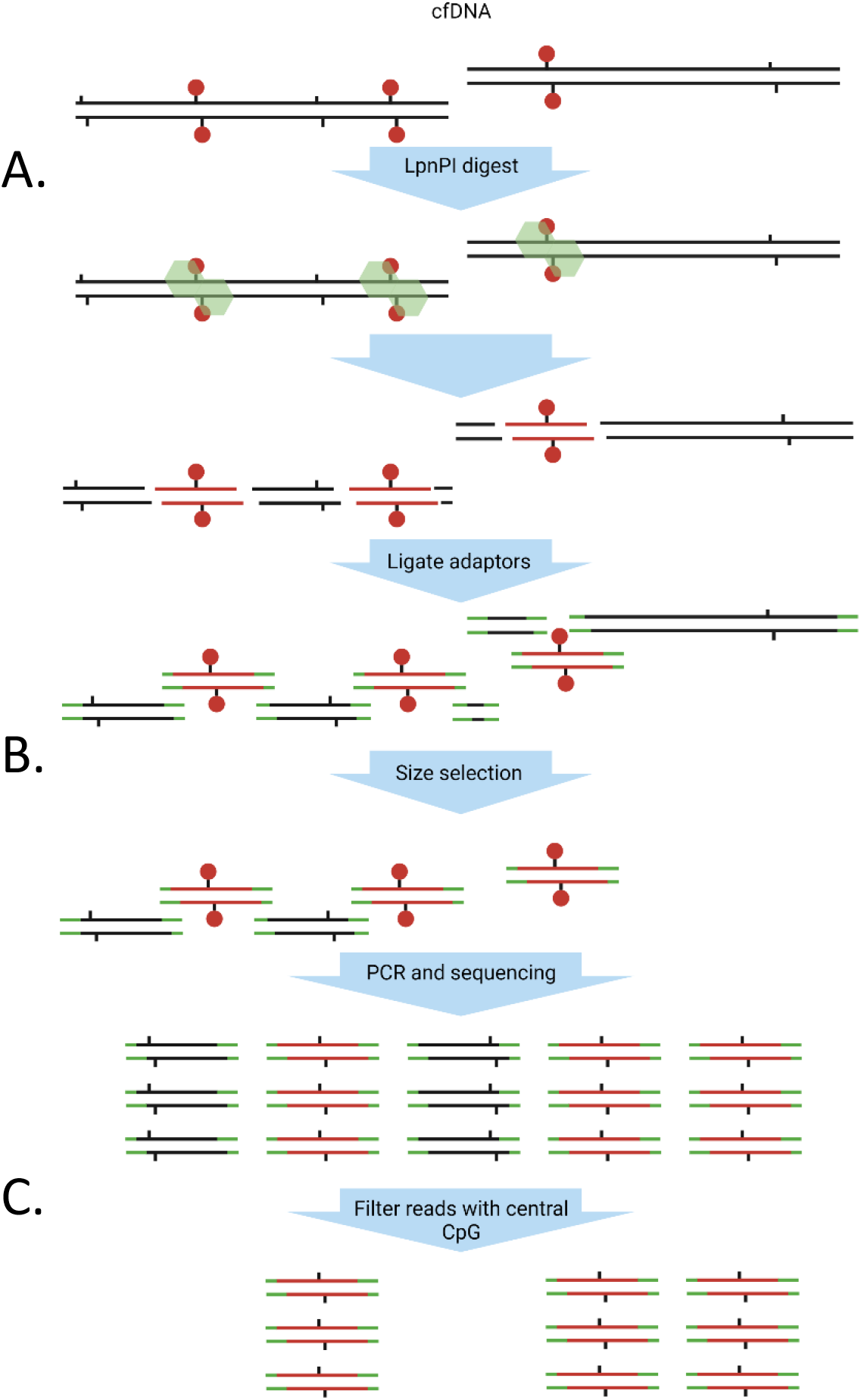
Schematic workflow of MeD-seq on cfDNA. **A**. The LpnPI endonuclease recognizes methylated CpG motifs and incises the DNA 16 bp upstream and downstream to generate 32bp fragments. **B**. After adaptor ligation, size fractionation is performed using the Pippin HT platform. **C**. All fragments of the selected size are amplified by PCR and sequenced. Only fragments with a CpG on the central position pass the filter and are considered as methylated reads. Green: LpnPI endonuclease, Black tabs: Cpg-site, Red circles: CpG methylation.

Here, we demonstrate the applicability of the MeD-seq method on cfDNA from plasma of healthy subjects, and metastatic cancer patients. We explore the compatibility of MeD-seq with various blood collection tubes, cfDNA isolation methods and cfDNA storage buffers and show that MeD-seq enables disease load monitoring in metastatic cancer patients.

## RESULTS

### Successful MeD-seq analysis from 10 ng cfDNA irrespective of blood collection tube

For successful MeD-seq analysis of cfDNA two crucial steps can be identified: **1)** the already fragmented cfDNA (∼150bp) needs to be properly digested in a methylation-dependent manner by the LpnPI enzyme into 32bp fragments, and **2)** the low cfDNA input needs to yield sufficient library-prepped DNA for subsequent sequencing. To ensure proper digestion in the first step, all resulting sequencing reads are filtered for the presence of an LpnPI restriction site at the expected position. This is particularly important for cfDNA since, in contrast to genomic DNA with a high molecular weight, undigested cfDNA is of short size and is therefore not necessarily excluded by the DNA fragment size selection step during library preparation.

We first evaluated the potential influence of the type of blood collection tube and cfDNA input amount on the MeD-seq assay performance. Blood from 3 patients (M4, M10, and M19) was collected in EDTA and CellSave tubes during the same blood draw. cfDNA was isolated from the resulting plasma using the manual QIAamp kit and eluted in AVE buffer (RNase-free water with 0.04% NaN_3_). Subsequently, for each sample both 10 ng of cfDNA and the maximal volume of 8 µl of cfDNA (>30 ng cfDNA) were used as input for MeD-seq. The cfDNA input in nanograms was kept equal per patient between EDTA and CellSave-derived cfDNA. The MeD-seq assay was successful regardless of input amount (10 ng vs >30ng). No differences were observed between CellSave and EDTA in the percentage of duplicate reads and the percentage of reads passing the LpnPI filter (Supplementary Figure S1). Principal component analysis of all obtained genome-wide methylation profiles showed that EDTA and CellSave-derived methylation profiles obtained from the same patient were clustered together, irrespective of input amount (Figure 2A and 2B).

**Figure 2.**
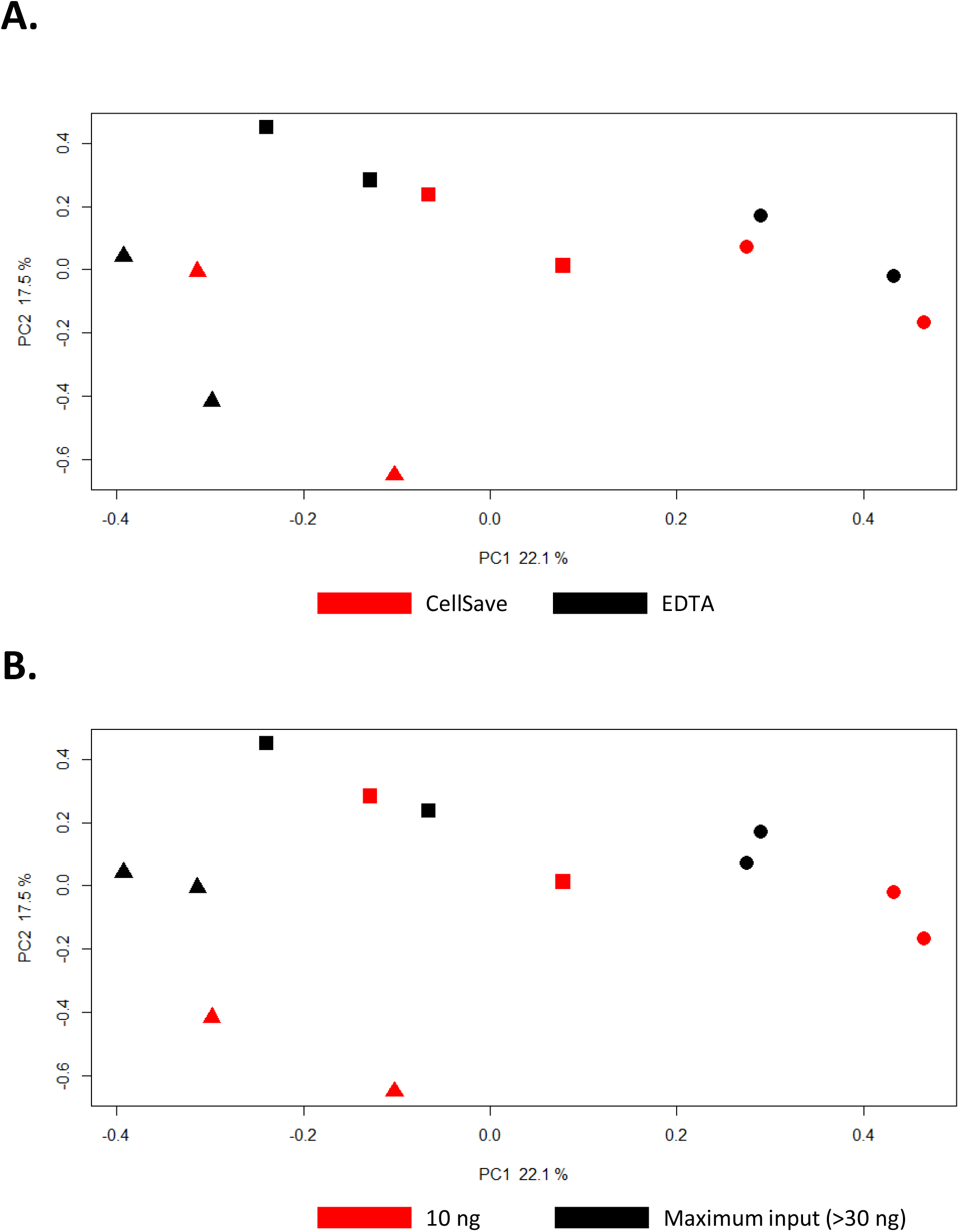
Principal Component Analysis of MeD-seq cfDNA methylation profiles using different blood tubes and input amounts. Principal components were calculated for cfDNA methylation profiles of 3 patients from which blood was collected at the same time in EDTA and CellSave tubes and either 10 ng cfDNA or the maximal input of 8 µl DNA was used. The maximal input amount was kept equal for EDTA and CellSave within 1 patient. PC1 and PC2, with the explained variances, are shown on the x-axis and y-axis respectively. Samples from the same patient are indicated by the shapes used, where each icon represents 1 cfDNA sample (patient M4 by ▲, patient M10 by ■, and patient M19 by ⍰). In **A**. samples are coloured based on the tube the blood was collected in (EDTA in black and CellSave in red). In **B**. samples are coloured based on the cfDNA input amount used, 10 g in black and maximum input in red (>30 ng).

In total, 51 QIAamp-isolated cfDNAs (both derived from patients and healthy blood donors) were then subjected to MeD-seq analysis, of which 4 had an input of less than 10 ng cfDNA, 34 had 10 ng cfDNA input, and 13 had more than 10 ng. For all samples with less than 10 ng input the yield after library preparation was insufficient to continue with sequencing, whereas both for samples with 10 ng input and more than 10 ng input only 1 sample failed at this stage (failure rate: 100%, 2.9%, and 7.7%, respectively; Chi-square p<0.001). From the foregoing we can conclude that the current MeD-seq protocol can be used to analyse plasma-derived cfDNA from both EDTA and CellSave blood collection tubes, provided at least 10 ng of cfDNA is available in a maximal volume of 8 µl.

To investigate the reproducibility of MeD-seq on cfDNA samples, we have calculated Pearson correlations between the cfDNA methylation profiles obtained from the 4 biological replicates of M4, M10, and M19 (different blood collection tubes and input amount) and compared these to the correlations for these samples with 9 healthy blood donors. As shown in Supplementary figure S2, observed correlations between biological replicates were significantly higher in all 3 patients compared to the correlations with healthy blood donors (Mann-Whitney U test, p < 0.005).

A selected panel of genes with leukocyte-specific methylation (17) showed consistent MeD-seq profiles in HBDs (median Pearson correlation = 0.85, range: 0.74 – 0.92, Supplementary figure S3), again demonstrating that MeD-seq yields reproducible results on cfDNA.

### Not all cfDNA isolation methods are compatible with MeD-seq analysis

In addition to the 45 QIAamp isolated cfDNA samples with at least 10 ng cfDNA input that were successfully analysed by MeD-seq, we also performed MeD-seq analysis on 37 cfDNA samples isolated by the semi-automated QiaSymphony platform. As shown in Figure 3A and B, cfDNA isolation using the semi-automated QiaSymphony platform (n=37) resulted in a significantly lower percentage of reads passing the LpnPI filter and duplicate reads compared to QIAamp isolated samples (n=45) (Mann-Whitney U test, p<0.001). In fact, for most QiaSymphony isolated samples the threshold of 10% LpnpI filtered reads was not met in the first 2M reads after which sequencing was aborted. The significant difference in filtered reads and duplicate reads isolated with QIAamp isolation and the QiaSymphony platform was shown using samples from different individuals. To exclude the possibility that our observations are cohort-specific instead of isolation method related, we isolated cfDNA from different aliquots of the same plasma sample (n=2) using different isolation methods, in which we also included the semi-automated Maxwell platform next to the already mentioned manual QIAamp kit and semi-automated QiaSymphony platform. The percentages of duplicate reads and reads passing the LpnPI filter of paired samples were also lower in the QiaSymphony sample for patient M69 compared to QIAamp and Maxwell (Figure 3C and 3D). For patient M12, the QiaSymphony isolation did not yield sufficient cfDNA for MeD-seq analysis, whereas both QIAamp and Maxwell generated comparable results to those obtained for patient M69. Pearson correlation analysis showed that the cfDNA methylation profiles obtained from paired QIAamp and Maxwell isolated samples correlated well (Pearson’s *r* >0.8), whereas the QiaSymphony isolated cfDNA profile was less well correlated (Pearson’s *r* = 0.62 compared with Maxwell isolation and *r* = 0.64 compared with QIAamp). Although the effect of QiaSymphony isolation compared to QiaAMP and Maxwell was only shown in 1 paired sample, together with our observations in the 82 unpaired cfDNAs (Figure 3A and 3B), results are indicative of an inhibited enzymatic activity of the LpnPI endonuclease in samples isolated by QiaSymphony. In cfDNA samples this results in sequencing of predominantly undigested cfDNA molecules not removed from the library preparation by size selection. Reads originating from these undigested molecules will be removed from the data by the LpnPI-sfilter, which only retains reads with a CpG dinucleotide at the central base pair position. Therefore, the low percentage of reads passing this filter in samples isolated on the QiaSymphony platform is characteristic for impaired LpnPI digestion.

**Figure 3.**
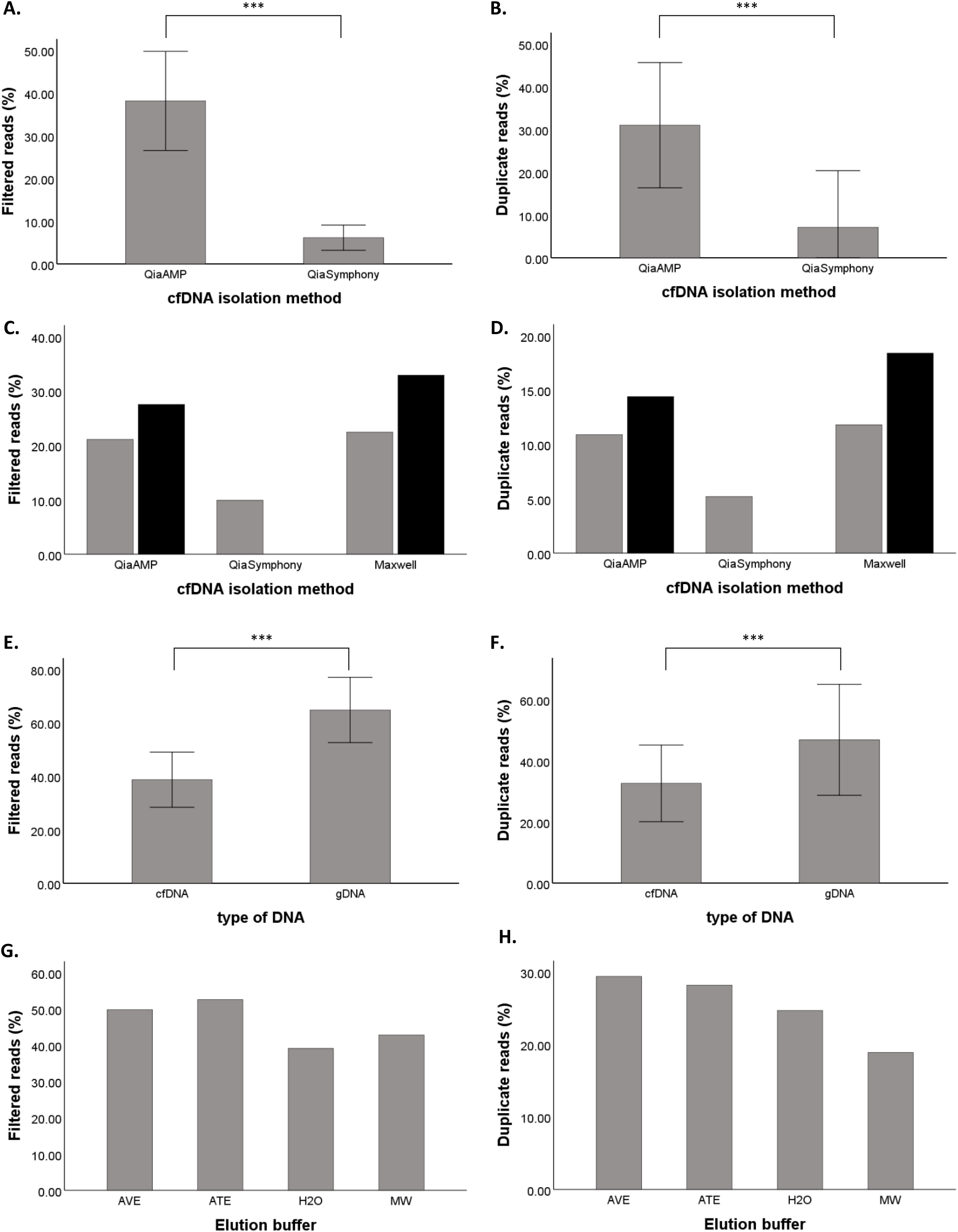
The compatibility of different isolation methods with MeD-seq analysis. In **A**. and **B**. percentages of LpnPI filtered reads and duplicate reads are shown for QIAamp (n=45) and QiaSymphony (n=37) isolated cfDNA samples. In **C**. and **D**. filtered reads and duplicate reads are shown of cfDNA isolated from aliquots of the same plasma from patients M12 (black) and M69 (grey) by QIAamp, QiaSymphony, and Maxwell. In **E**. and **F**. a significantly lower percentage of filtered and duplicate reads is shown for cfDNA compared to gDNA samples. In **G**. and **H**. filtered reads and duplicate reads are shown for genomic MCF7 DNA in the different elution buffers. ***: P ≤ 0.001, AVE buffer (QIAamp): RNase-free water with 0.04% NaN3; ATE buffer (QiaSymphony): 10 mM Tris-HCl pH 8.3, 0.1 mM EDTA, and 0.04% NaN3; MW buffer (Maxwell): 10 mM Tris, 0.1 mM EDTA, pH 9.0.

As these cfDNA isolation kits contain different elution buffers, we subsequently compared the performance of MeD-seq on 10 ng genomic DNA isolated from MCF7 cells by QIAamp, but eluted in the different elution buffers or water. In general, genomic MCF7 DNA yielded more reads passing the LpnPI-filter as well as a higher percentage of duplicate reads compared to cfDNA samples (Mann-Whitney U test, p <0.001, Figure 3E and 3F). This is likely due to the fact that the DNA fragment size selection step in the protocol is unable to eliminate undigested small cfDNA molecules from the sequencing library. Irrespective of the elution buffer used, the obtained percentages of duplicate and filtered reads were sufficient and comparable to water, and the MeD-seq data obtained from the different MCF-7 samples were highly correlated (Pearson’s *r*>0.90), indicating that differences in elution buffers are not causing the observed LpnPI inhibition in QiaSymphony isolated samples (Figure 3G and 3H).

### Vacuum concentration of cfDNA samples is compatible with the MeD-seq assay

In the current protocol the maximal sample input volume is 8 µl, requiring a cfDNA concentration of at least 1.25 ng/µl to enable the minimally required input of 10 ng cfDNA. Although cfDNA yields are dependent on tumour type, disease stage, moment of blood draw, and isolation method, in our hands the obtained cfDNA concentrations for the majority of 2 ml plasma samples are below this threshold even though the total amount of 10 ng is available. To increase the number of cfDNA samples compatible with MeD-seq analysis, we therefore evaluated whether it was possible to apply vacuum concentration to cfDNA samples before MeD-seq. As a first step we checked whether the increased salt concentrations after vacuum concentration of the samples influenced the enzymatic activity of the LpnPI restriction enzyme. For this purpose, we eluted genomic DNA from MCF7 cells in either AVE buffer (QIAamp kit), MW buffer (Maxwell kit) or water. Subsequently, genomic MCF7 DNA was diluted to 10 ng in 20 µl of the respective elution buffer and the volume was reduced by vacuum concentration to 8 µl. Unlike cfDNA, which is already fragmented, undigested genomic DNA is excluded from the library preparation due to the DNA fragment size selection step included in the protocol. Therefore, in case of incomplete digestion by LpnPI, for genomic DNA only the small fraction of digested genomic DNA molecules will be subjected to sequencing, whereas for cfDNA undigested cfDNA molecules are included in the sequencing library as well. Based on this information, incomplete digestion of genomic DNA is expected to result in increased numbers of duplicate reads passing the LpnPI-filter, whereas incomplete digestion of cfDNA results in lower numbers of reads passing the LpnPI-filter (Figure 3C and 3D). As shown in Figure 4A and 4B, vacuum concentration resulted in a higher percentage of both filtered reads and duplicate reads with MW buffer compared to both AVE buffer and water. As explained above these results suggest incomplete digestion of the genomic DNA by LpnPI in MW buffer. MW buffer contains EDTA, which is known to inhibit metallo-enzymes by chelation of the metal ion needed for their catalytic activity (18). These results suggest that for genomic DNA in water or EDTA-free buffers it is possible to vacuum concentrate samples to achieve a minimal input of 10 ng. To subsequently test whether this is also true for cfDNA, we used 3 plasma aliquots from the same metastatic breast cancer patient sample (P71), isolated cfDNA by QIAamp and eluted the cfDNA in AVE buffer, MW buffer or water. For each buffer 10 ng of cfDNA was either analysed by MeD-seq directly or first diluted to 20 µl and vacuum concentrated to 8 µl before MeD-seq analysis. As expected, we again observed a clear indication that the enzymatic activity of LpnPI was inhibited by vacuum concentration of the MW cfDNA sample, as the percentage of reads passing the LpnPI filter was reduced (Figure 4C). Vacuum concentration of cfDNA in AVE buffer or water did not alter the percentage of LpnPI-filtered reads or duplicate reads compared to their non-concentrated control (Figure 4C and 4D). Pearson correlations between samples with and without vacuum concentration were 0.73 for cfDNA in AVE buffer, 0.72 for cfDNA in water and only 0.48 for cfDNA in MW buffer (Supplementary Table 1Bxs). Similarly, principal component analysis showed two distinct clusters for P71 and MCF7 containing all non-concentrated samples as well as the concentrated samples in water and AVE buffer, whereas the concentrated samples in MW buffer were outliers for both genomic MCF7 DNA and cfDNA from patient P71 (Figure 4E). Together these results suggest vacuum-concentration is compatible with MeD-seq analysis but may be hindered by the presence of increased EDTA concentrations in the buffer after concentration.

**Table 1:**
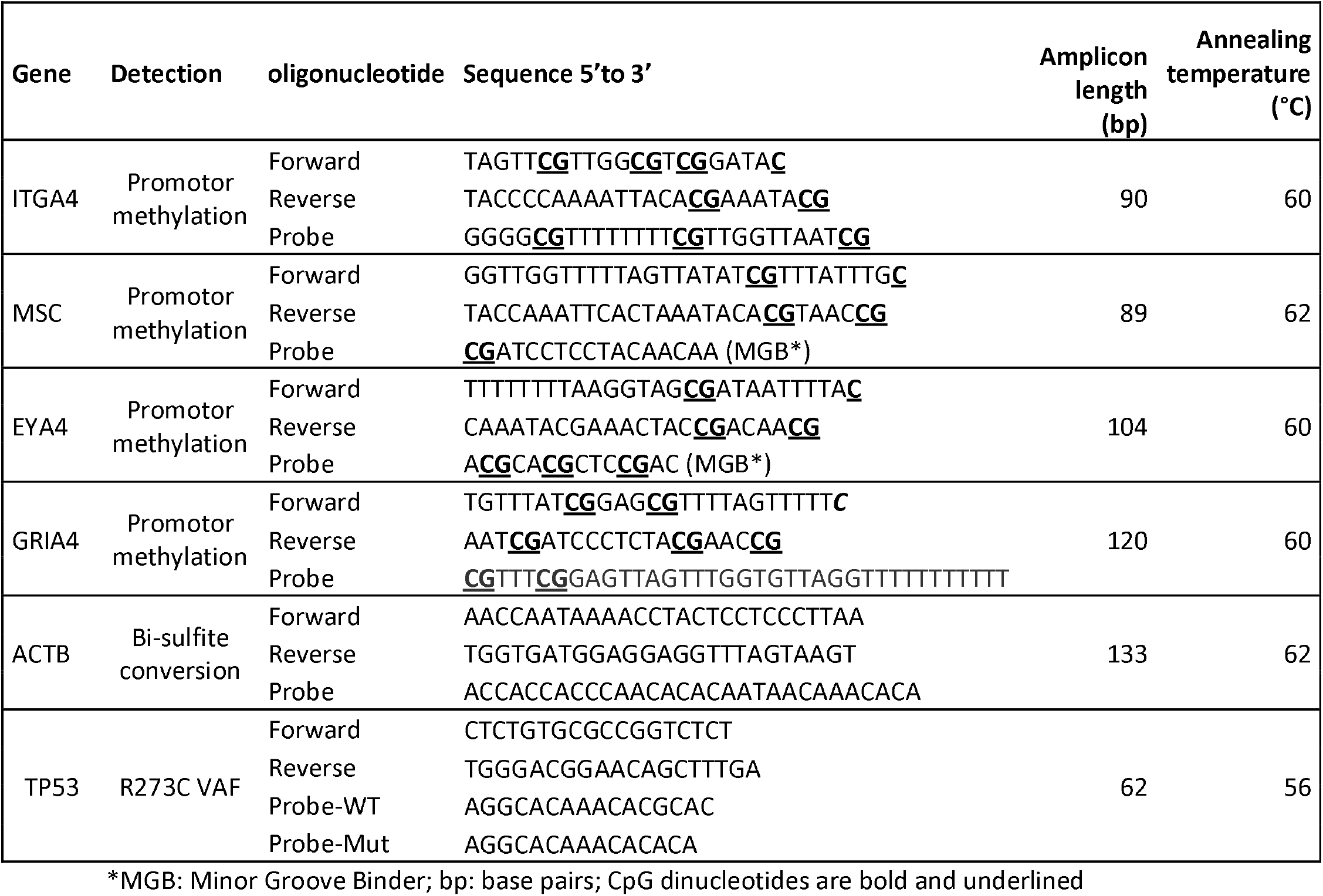
Primer and probe sequences. CpG dinucleotides are bold and underlined.

**Figure 4.**
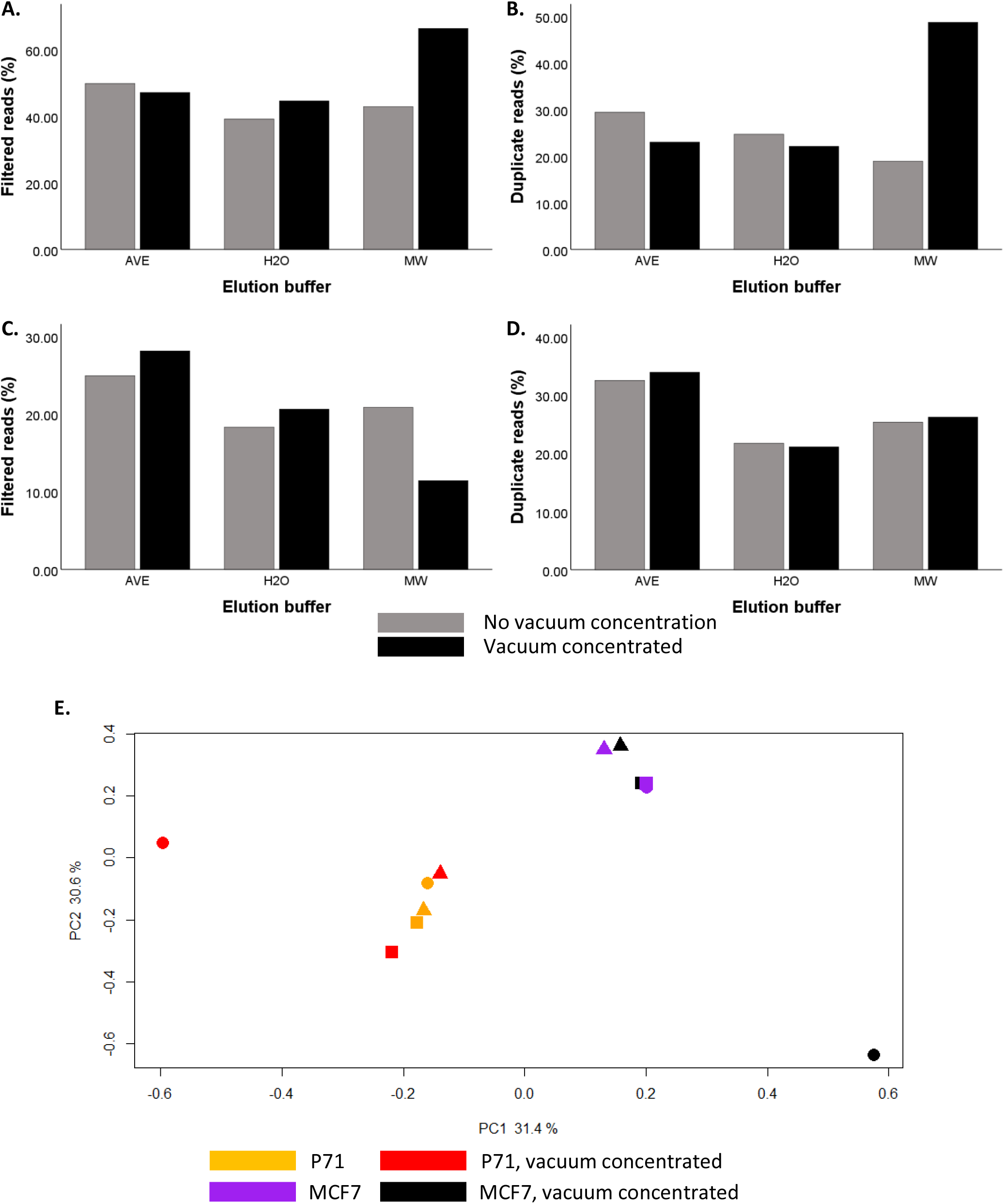
The compatibility of vacuum concentrated samples in different buffers with MeD-seq analysis. The top two panels show results for 10 ng of genomic MCF7 DNA dissolved in AVE buffer, water, or Maxwell elution buffer analysed by MeD-seq with and without preceding vacuum concentration of the sample. In **A**. the percentage of LpnpI filtered reads and in **B**. the percentage of duplicate reads is shown for the different elution buffers with (in black) and without (in grey) vacuum concentration. The middle two panels show results for different aliquots of 10 ng cfDNA from the same plasma sample dissolved in AVE buffer, water, or Maxwell elution buffer analysed by MeD-seq with and without preceding vacuum concentration of the sample. In **C**. the percentage of LpnpI filtered reads and in **D**. the percentage of duplicate reads is shown for the different elution buffers with (in black) and without (in grey) vacuum concentration. **E**. Principal components were calculated for cfDNA methylation profiles of genomic MCF7 DNA and 3 plasma aliquots from a single metastatic breast cancer patient dissolved in AVE (■), H2O (▲), MW (⍰) buffer with and without vacuum concentration. PC1 and PC2, with the explained variances, are shown on the x-axis and y-axis respectively. Each icon represents 1 cfDNA sample: samples coloured in red and black were vacuum-concentrated whereas samples in orange and purple were not.

### MeD-Seq for disease load monitoring in CRLM patients before and after surgery

To evaluate whether genome-wide cfDNA methylation profiles are representative of tumour load, we performed MeD-seq analysis on pre- and post-operative cfDNA samples from 8 patients with operable colorectal liver metastases (CRLM). For comparison, profiles of 9 healthy blood donors (HBDs) were included as well. Seven out of 8 pre-operative samples were also analysed for KRAS, TP53, or PIK3CA mutations by digital PCR, which were found in all 7 samples with variant allele frequencies (VAFs) of 30.2% (M1), 10.9% (M2), 27.3% (M4), 67.3% (M8), 12.9% (M10), 19.7% (M19), and 41.4% (M34) verifying the presence of ctDNA in these samples.

Principal component analysis (PCA) as well as unsupervised hierarchical clustering analysis showed that pre-operative CRLM samples were very distinct from HBDs, whereas 5 days after surgery methylation profiles were more comparable to HBDs (Figure 5A and 5B). As is shown in Figure 5C, the summarized Z-score based on all regions in all pre-surgical samples is clearly elevated (average: 31.52; range: 10.24 - 69.48) compared to HBDs (average: 0.00; range: -0.95 - 1.82), whereas the summarized Z-score in all post-surgical samples (average: 0.06; range: -1.47 - 2.91) is in the same range as those from HBDs. The summarized Z-scores before surgery were strongly correlated with the observed VAFs for KRAS, TP53, or PIK3CA mutations in the same samples (Pearson *r* = 0.90, p = 0.006), indicating the summarized genome-wide deviation in methylation profile reflects the level of ctDNA within the total pool of cfDNA.

**Figure 5.**
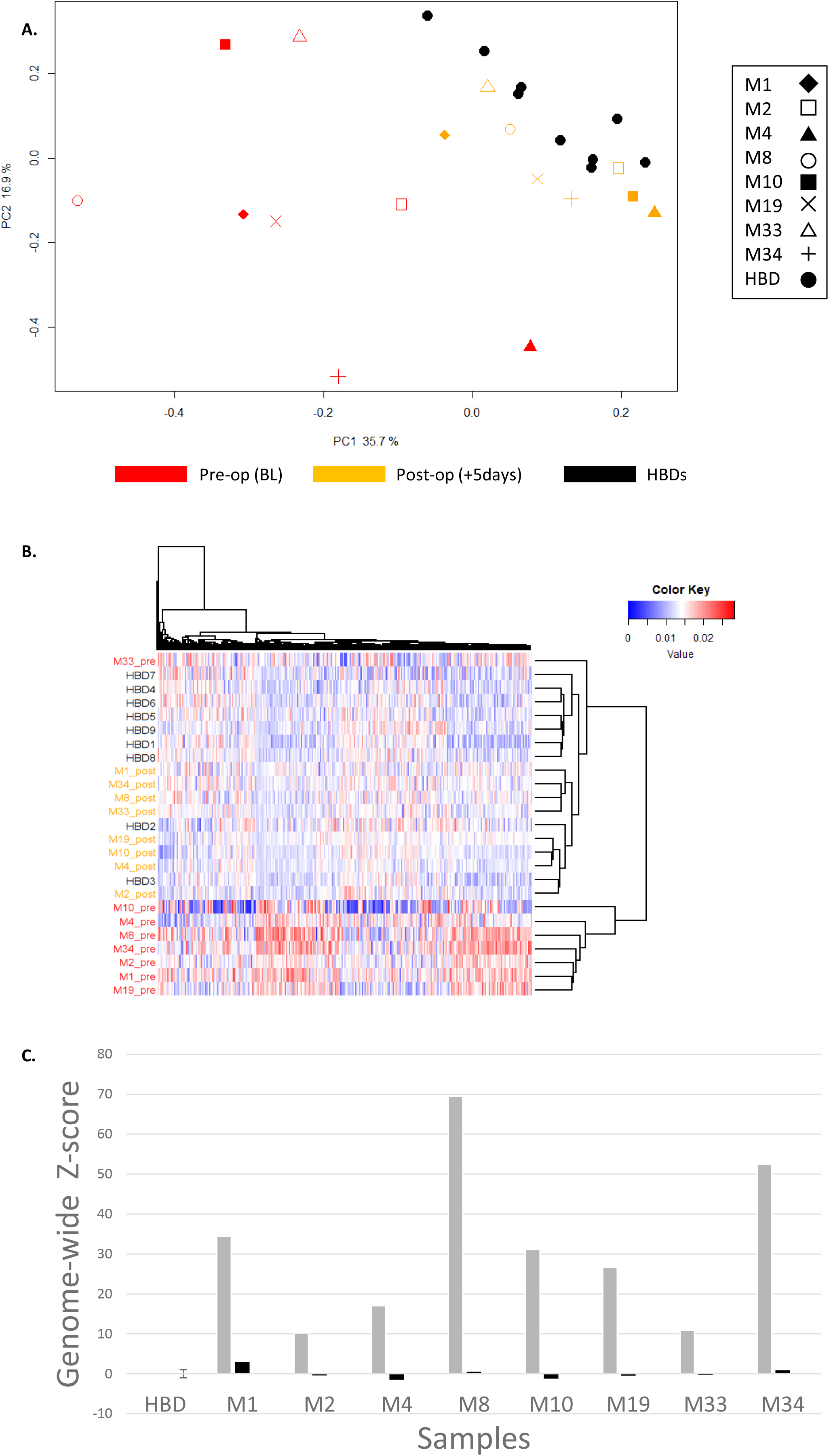
Comparision of cfDNA methylation profiles from HBDs and CRLM patients before and after surgery. **A**. Principal components were calculated for cfDNA methylation profiles of 9 HBDs (4 males and 5 females; in black) and a pre-(BL; in red) and postoperative (+5d; in orange) sample of 8 CRLM patients. PC1 and PC2, with the explained variances, are shown on the x-axis and y-axis respectively. Shapes represent patients, where each icon represents 1 cfDNA sample. **B**. Unsupervised hierarchical clustering of methylation profiles again shows a distinct cluster for pre-operative CRLM samples (red). HBDs (black) and post-operative CRLM samples (orange) show more similar methylation profiles. **C**. Obtained methylation profiles were summarized in a genome-wide Z-score as an overall measure for the deviation of the sample from the average normal methylation profile in HBDs. Pre-operative genome-wide Z-scores are depicted in grey and post-operative genome-wide Z-scores in black.

Multiple methylation markers for (metastatic) CRC disease monitoring have already been described in literature, including ITGA4, MSC, EYA4, GRIA4, MAP3K14-AS1 (19), B4GALT1 (20), BCAT1, IKZF1 (21); (22), SEPT9, and SHOX2 (23); (24). MeD-seq results are shown for these markers in paired pre- and post-operative CRLM blood samples for patients M1, M2, M4, M8, M10, M19, M33, and M34, as well as for the corresponding CRLM tissues for patients M4, M10, and M19. (Figure 6A & Supplementary Figure S4). Distinct differences between methylation profiles were observed in pre-operative blood samples (either with available tissue samples) on one side and the post-operative blood samples on the other side for ITGA4, in 7 patients; MSC, in 6 patients; GRIA4, in 6 patients; EYA4, in 7 patients; BCAT1, in 7 patients; SEPT9, in 8 patients; and IKZF1, in 4 patients. In contrast, SHOX2, B4GALT1, and MAP3K14-AS1 showed either similar methylation levels in all samples (SHOX2) or virtually no methylation in any of the samples analysed (B4GALT1 and MAP3K14-AS1). We validated results for ITGA4, MSC, GRIA4, and EYA4 in paired pre- and post-operative blood samples from 10 CRLM patients, 8 of which were also analysed by MeDseq, using an independent method (qMSP). ITGA4, MSC, and EYA4 showed a statistically significant decrease in CRLM patients 5 days after surgery compared to the paired baseline sample prior to surgery, whereas the HBD controls do not show any methylation for these markers. GRIA4 also showed a decrease in the same samples trending towards significance (exact sign test p=0.07) (Figure 6B). Subsequent Pearson correlation analysis for those samples analysed both by MeD-seq and qMSP showed that for EYA4, ITGA4, and MSC results from both methods were significantly correlated (EYA4 *r* = 0.78, p=0.001; MSC *r* = 0.92, p<0.0001; and ITGA4 *r* = 0.59, p=0.026). MeD-seq and qMSP results for GRIA4 again showed the same trend with borderline significance (r = 0.48, p=0.08, Supplementary figure S4). Together, these results support the reliability of the MeD-seq assay on cfDNA samples.

**Figure 6.**
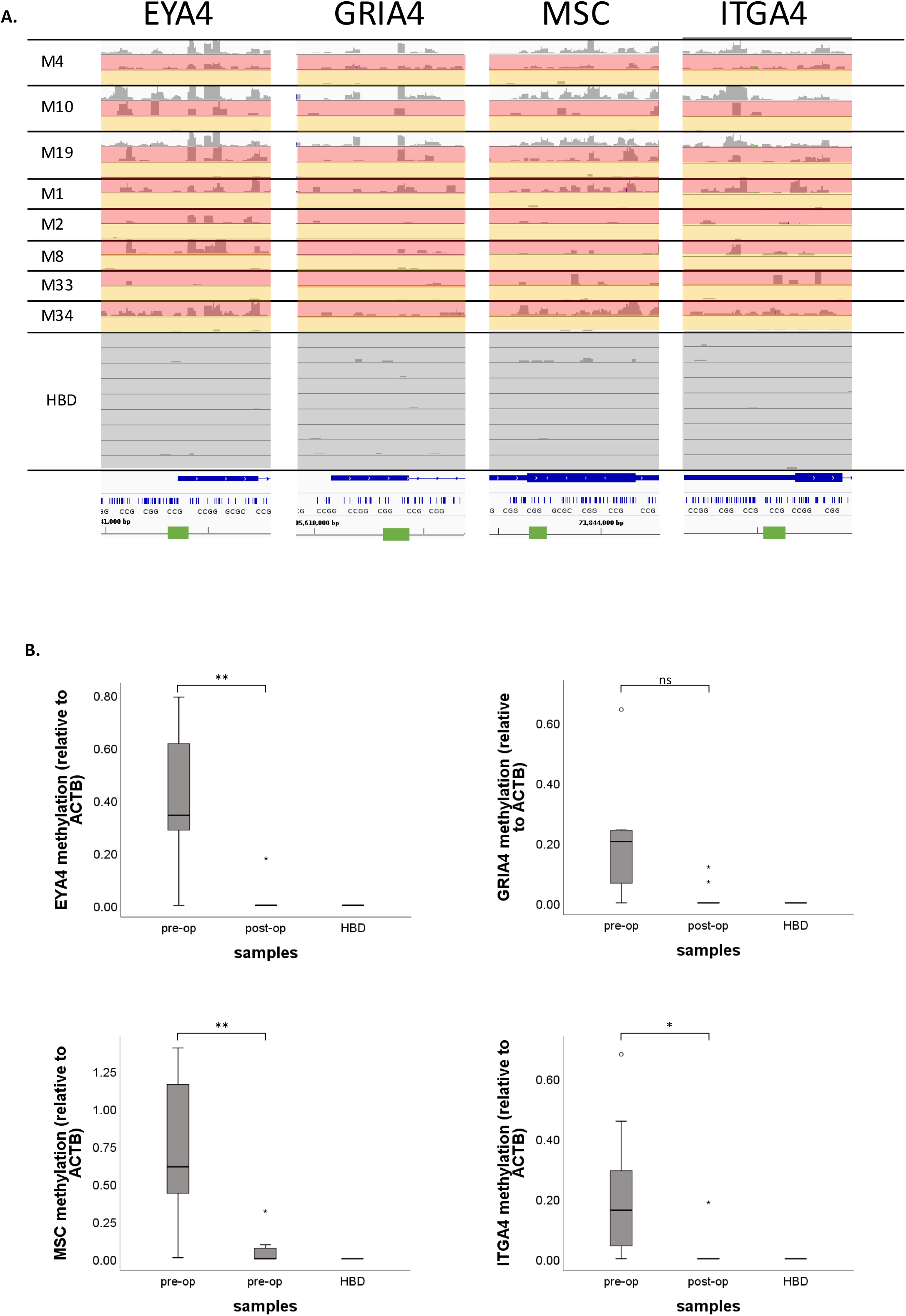
MeD-seq results for published biomarkers on cfDNA from HBDs and paired tissue, pre-operative and post-operative cfDNA from CRLM patients. **A**. Sequenced MeD-seq reads for EYA4, GRIA4, MSC, and ITGA4 of CRLM tissue (white), pre-operative cfDNA (red), and post-operative cfDNA (orange) from CRLM-patients and HBDs (black). Results are visualized in Integrative Genomics Viewer (IGV) v.2.9.4. The green bars indicate the locations of the respective qMSP amplicons. **B**. qMSP-based methylation levels relative to the ACTB reference gene are shown for EYA4 (top-left), GRIA4 (top-right), MSC (bottom-left), and ITGA4 (bottom-right) at baseline (BL) and 5 days post-surgery (+5d) in cfDNA of 10 CRLM patients and 3 healthy blood donors (HBDs). A median decrease in methylation levels was observed for EYA4, GRIA4, MSC, and ITGA4 5 days after surgery compared to BL in CRLM patients (exact sign test; p=0.004, p=0.07, p=0.002, and p=0.039, respectively).

## DISCUSSION

This study demonstrates the feasibility of genome-wide cfDNA methylation profiling on liquid biopsies using the recently developed MeD-seq assay (16). Compatibility with different blood collection tubes, cfDNA isolation platforms, and vacuum-concentration of samples was explored to facilitate the future implementation of this method by other laboratories.

To further substantiate its potential for real-time tumour load monitoring, we applied the MeD-seq assay to paired liquid biopsies of 8 patients with colorectal liver metastases before and after surgery. We related methylation values of all regions with sufficient data to their methylation status in HBDs to come to a single summarized Z-score as was described before for chromosomal aneuploidy (25). This score represents the global deviation in methylation in a patient cfDNA sample compared to a reference pool of normal cfDNAs. The potential value for this type of scoring was nicely illustrated by the high correlation between this score and the observed VAFs for known mutations before surgery as well as by the sharp decrease in genome-wide Z-scores in our 8 CRLM patients after surgery. Together these observations indicate that the global deviation in methylation profile reflects the amount of ctDNA within the pool of cfDNA.

Next to the MeD-seq we describe here, cfMeDIP-seq represents another promising method for cfDNA methylation profiling (26). With the current protocol, a similar number of reads (30-35M) is estimated to be necessary for both techniques. A major advantage of the cfMeDIP-Seq is that it only requires 1 ng cfDNA input, whereas the current MeD-seq protocol requires 10 ng. Input below 10 ng cfDNA invariably resulted in insufficient library yield in our hands (n=4). However, for most 2 ml plasma aliquots a yield of 10 ng cfDNA is feasible, especially given the potential compatibility of subsequent vacuum concentration with MeD-seq analysis we observed in a small number of samples.

Compared to cfMeDIP-seq (14), the MeD-Seq protocol requires less hands-on time and is therefore more straightforward. For MeD-seq, cfDNA samples are digested overnight, used as input for standard Illumina library preparation, size-selected and sequenced. For cfMeDIP-seq on the other hand, one needs to additionally prepare filler DNA, and purify and perform quality control on the sample after immunoprecipitation. The number of required reads for MeD-seq depends on the number of reads passing the LpnPI filter, which is lower in fragmented cfDNA compared to genomic DNA due to the presence of undigested cfDNA fragments in the same size range as the digested fragments. In that respect, selection of digested cfDNA from undigested cfDNA as incorporated in the recently described cfDNA-RRBS protocol may also represent a valuable addition to our current MeD-seq protocol to reduce the number of background reads (12, 13)). Although we did not perform a head-to-head comparison, theoretically MeD-seq is able to give a higher and less biased overall coverage of CpG dinucleotides (15, 16).

In our view MeD-seq analysis of cfDNA has 2 distinct potential applications: 1) discovery of relevant DMRs for the development of subsequent marker panels, and 2) direct use as diagnostic assay. In literature so far 10 methylation markers have been described for disease load monitoring and/or detection of minimal residual disease in liquid biopsies of CRC patients, namely EYA4, GRIA4, ITGA4, MAP3K14-AS1, MSC ((19)), SEPT9, SHOX2 ((23)), BCAT1, IKFZ1 (21) (22), and B4GALT1 (20). With MeD-seq, we detected differentially methylated regions between pre- and post-operative samples for EYA4, GRIA4, ITGA4, and MSC, which we validated with qMSP. Furthermore, MeD-seq detected differentially methylated regions for BCAT1, SEPT9 and IKZF1 but not for B4GALT1, MAP3K14-AS1 and SHOX2.

Although we only had 8 paired patient samples in our proof of principle experiment, the MeD-seq assay can be used for direct discovery screens on cfDNA instead of tissues or cell lines to identify additional, perhaps even more relevant, markers for liquid biopsy-based disease monitoring in CRC patients.

Next to discovery of disease-specific biomarkers in cfDNA, one could also envision the direct clinical implementation of the MeD-seq approach to generate genome-wide cfDNA methylation profiles for patients over time. Although this is more expensive than the application of targeted panels by PCR-based methods, costs for cfDNA MeD-seq are comparable to that of targeted cfDNA mutation panels as well as to costs associated with a standard CT-scan of the thorax and abdomen. Results in literature so far show that cfDNA methylation profiling may show an increased overall sensitivity for disease detection, longitudinal disease monitoring, and tumour classification(26); (10).

## CONLUSIONS

In conclusion, we here present a novel method for cfDNA methylation profiling and show that this method is compatible with 10 ng of cfDNA isolated manually by QIAamp or semi-automated by Maxwell from plasma obtained from EDTA or CellSave blood collection tubes. The potential of the MeD-seq assay is shown by the obtained paired cfDNA methylation profiles in CRLM patients before and after surgery and their comparison to cfDNA methylation profiles in HBDs. Our assay provides a suitable tool both for the discovery of methylation markers in cfDNA samples and direct monitoring of tumour load in metastatic cancer patients.

## MATERIALS & METHODS

### Cell lines and patient samples

For optimization experiments genomic DNA from MCF7 cells was used, which were obtained from the American Type Culture Collection (ATCC, Manassas, VA). MCF7 cells were cultured in Gibco™ RPMI 1640 glutamax (Thermo Fisher Scientific, Waltham, MA) supplemented with 10% heat-inactivated fetal bovine serum, 80µg/ml streptomycin, and 100µg/ml penicillin G.

Coded metastatic cancer patient samples were obtained from various clinical studies, all of which were approved by the Medical Ethics Committee from the Erasmus University Medical Center (MEC 17-238, MEC 15-289, and MEC 16-499). In short, 10 ml of blood was collected from patients in EDTA tubes, stabilizing CellSave tubes (Menarini Silicon Biosystems, Castel Magiore, Italy), or simultaneously in both tube types. In addition, 10 ml of blood was obtained from 9 consenting anonymous healthy blood donors (HBDs; 5 females and 4 males) via the Dutch National blood bank (Sanquin) in CellSave tubes. Plasma was isolated from the obtained blood within 24 hours (EDTA) or 96 hours (CellSave) after blood draw by 2 sequential centrifugation steps at room temperature (10 minutes at 1711g followed by 10 minutes at 12000g) and stored at −80 °C (27).

### (cf)DNA isolations and quantification

Genomic DNA from MCF7 cells was isolated by proteinase K digestion at 65°C for 30 minutes followed by purification using the QIAamp circulating nucleic acid kit (Qiagen, Venlo, The Netherlands). Genomic DNA from frozen tissue sections of colorectal liver metastases was isolated using the NucleoSpin Tissue kit according to the manufacturer’s guidelines. cfDNA was isolated from 2 ml of plasma using either the manual QIAamp circulating nucleic acid kit (Qiagen), or the semi-automated QIAsymphony DSP Circulating DNA Kit (Qiagen) and Maxwell® RSC ccfDNA Plasma Kit (Promega, Leiden, the Netherlands). DNA was eluted in elution buffers provided by the kits used (QIAamp kit: AVE buffer (RNase-free water with 0.04% NaN_3_), QiaSymphony kit: ATE buffer (10 mM Tris-HCl pH 8.3, 0.1 mM EDTA, and 0.04% NaN3), Maxwell buffer (10mM Tris, 0.1mM EDTA, pH 9.0)) or RNAse-free water as specified.

### MeD-seq assay

MeD-seq assays were essentially performed as previously described (16). In short, 8 µl genomic DNA (input ranged from 117-1728 ng) from frozen tissues, specified amounts of MCF7 genomic DNA or plasma-derived cfDNA were digested with LpnPI (New England Biolabs, Ipswich, MA) yielding 32 bp fragments around the fully methylated recognition site containing a CpG. Samples were prepped for sequencing using the ThruPLEX DNA-seq 96D kit (Rubicon Genomics, Takara Bio Europe, Saint-Germain-en-Laye, France) and purified on a Pippin HT system with 3% agarose gel cassettes (Sage Science, Beverly, MA). Libraries were multiplexed and sequenced on an Illumina HiSeq 2500 for 50 bp single reads according to the manufacturer’s instructions (Illumina, San Diego, CA). Samples were first sequenced until ∼2M reads and continued to a total of ∼20M reads only when the fraction of reads that passed the LpnPI filter (explained below) was at least 20%.

### Quantitative Methylation Specific PCR (qMSP)

The DNA methylation status of the CpG-island containing promoter regions of MSC, ITGA4, GRIA4, and EYA4 was determined by qMSP analysis on sodium bisulfite-treated cfDNA obtained before and 5 days after surgery of 18 CRLM patients as well as 3 HBDs. In brief, 5 ng cfDNA was modified using the EZ DNA Methylation kit (Zymo Research, Orange, CA), which induces chemical conversion of unmethylated cytosines into uracils. Specific primers were designed to amplify the methylated DNA sequence of all 3 promoter regions and resulting amplicons were quantified using TaqMan probes (Table 1). In addition, the modified, unmethylated sequence of the housekeeping gene β-actin (ACTB) was amplified as a reference (28). ACTB and MSC reactions were combined into a duplex reaction as were ITGA4 and GRIA4 reactions, whereas EYA4 reactions were performed separately. qMSP reactions were carried out in a 12.5 μl reaction volume containing 3µl of bisulfite-treated cfDNA, 300 nM (ACTB, MSC, ITGA4, GRIA4) or 500 nM (EYA4) of each primer, 250 nM probe, 6.25 µl 2x EpiTect MethyLight Master Mix (w/o ROX), and 1µl 50x ROX Dye Solution using the Mx3000P and Mx3005P QPCR Systems (Stratagene, La Jolla, CA). Only samples with a Ct for ACTB below 32 were included for data analysis to ensure sufficient DNA quality and quantity, resulting in paired data for 10 CRLM patients. Methylation values of the 3 target regions were normalized to the reference gene ACTB using the comparative Ct method (2-ΔCT) (29) and subsequently square root transformed to reduce skewness in the data distribution.

### Digital PCR (dPCR)

The presence of KRAS, TP53, and PIK3CA mutations in pre-surgical cfDNA of 7 CRLM patients was evaluated using digital PCR (dPCR). For KRAS, variant allele frequencies were determined by KRAS G12 and G13 mutation specific assays (Thermo Fisher Scientific) for patients M2, M4 and M19, whereas a KRAS mutation screening kit (Biorad, Hercules, CA) was applied to patient M10. The KRAS genotype was determined either directly in 7.5 µl cfDNA (patient M4) or after pre-amplification of 2 µl cfDNA (patients M2, M10, and M19). For TP53 R273 a mutation specific assay (Table 1) was designed to determine the variant allele frequencies in patients M1 and M8. For PIK3CA, the variant allele frequency was determined by the PIK3CA H1047 mutation specific assay (Thermo Fisher Scientific) for patient M34. The dPCR-analyses were performed either on a QuantStudio 3D dPCR system (Thermo Fisher Scientific) or a Naica System (Stilla Technologies, Villejuif, France). Pre-amplification and QuantStudio dPCR analyses were performed as described previously (30). Regarding the Naica system, up to 8 µl (pre-amplified) cfDNA was loaded onto the chip in the presence of a final concentration of 1x Perfecta MultiPlex ToughMix (QuantaBio, Beverly, MA, USA), 100 nM Fluorescein (VWR, Leicestershire, UK) and 1x FAM [mutant]/VIC [wild-type] labelled KRAS probe assays. After partitioning the sample into 30,000 crystals, the DNA copies were amplified in 45 PCR cycles (95°C for 10 minutes, followed by 45 cycles of 95°C for 30 seconds and 52 or 58°C for 15 seconds). After PCR, the acquired fluorescence data were analyzed with the CrystalMiner software version 1.6 (Stilla Technologies).

### Data processing

Dual indexed samples were demultiplexed using bcl2fastq software (Illumina). Subsequent data processing was carried out using specifically created scripts in Python, which include a trimming step to remove the Illumina adapters and a filtering step based on LpnPI restriction site occurrence between 13 and 17 bp from the 5’- or 3’ end of the read, after which DNA fragments which were methylated remain (Figure 1) (16). Reads passing the filter were mapped to the genome using Bowtie 2 (31). Using all unambiguously mapped reads, count scores were assigned to each individual LpnPI site in the genome. Outcome measures for a technically successful MeD-seq analysis include the following: 1) total number of obtained reads, 2) fraction of reads passing the LpnPI filter, and 3) the fraction of duplicate reads.

Subsequently, count scores for individual CpG sites were summarized into 2 kilobase (kb) regions surrounding all known transcription start sites (TSS) annotated in ENSEMBL, resulting in 57278 regions located on the autosomal chromosomes. After normalization (counts per million) using the total number of reads passing the LpnPI filter per sample, square root transformation was applied to reduce skewness in the data distribution. Regions with 0 counts in more than 25% of samples were removed, resulting in 39386 regions for CRLM samples (9 HBDs; 8 pre-surgical cfDNAs samples from CRLM patients) and 38879 regions for breast cancer samples (9 HBDs; 3 pre-vacuum samples eluted in AVE, H2O, and MW-buffer, 4 MCF7 samples) for further data analysis.

### Data analysis and statistical testing

#### Principal component analyses

Principal component analysis (PCA) was performed on the 50% most variable regions to reduce the dimensionality of the data. The normalized and transformed data were mean-centered and subsequently reduced to two principal components using the Singular Value Decomposition function (svd) in R v.3.6.3.

#### Genome-wide Z-score calculation

To generate one overall score for aberrant cfDNA methylation per patient sample, Z-scores were first calculated per region by subtracting the mean and dividing by the standard deviation of that respective region in a panel of nine healthy controls (HBDs). Resulting Z-scores per region were squared and summed for both patient samples and HBDs to get a genome-wide Z-score as described previously (25). For HBDs genome-wide Z-scores were calculated using a leave-one-out approach, in which one sample was compared to the remaining healthy controls. Calculations were performed in R v3.6.3 (www.R-project.org).

#### Additional analyses

Technical outcome measures from the MeD-seq assay were compared between groups using the chi-square or Mann-Whitney U test depending on the type of variable. Paired pre- and post-surgical qMSP data were analysed using the paired-samples sign test. These analyses were performed in IBM SPSS Statistics 25 and two-sided p values <0.05 were considered statistically significant. Pearson correlation analyses between DNA methylation profiles were performed using R v3.6.3 (www.R-project.org).

## DECLARATIONS

### Ethics approval and consent to participate

Coded metastatic cancer patient samples were obtained from various clinical studies (MEC 17-238, MEC 15-289, and MEC 16-499), all of which were approved by the Medical Ethics Committee from the Erasmus University Medical Center. All patients signed an informed consent.

### Consent for publication

Not applicable

### Availability of data and materials

The dataset supporting the conclusions of this article is available in the NCBI Sequence Read Archive (SRA; https://www.ncbi.nlm.nih.gov/sra) under accession number SUB9929930.

### Authors’ contributions

TD, RB, VdW, ZA, WvIJ, and MMJvdP performed the described experiments. JB, TD, RB, WvIJ and SW were involved in bioinformatical and statistical data analysis. TD, RB, JG, JWMM, and SW wrote the manuscript. LA, DJG, LFvD, MPJKL, CV, and SS conducted the clinical studies and made samples available to this study. LA, SS, CV, JWMM, JG, and SW conceived the study and obtained the necessary funds for execution of the study.

## Acknowledgements

We highly appreciate the help of Jean Helmijr and Maurice Jansen with the digital PCR analyses.

## Figure legends

**Supplementary Figure S1.**
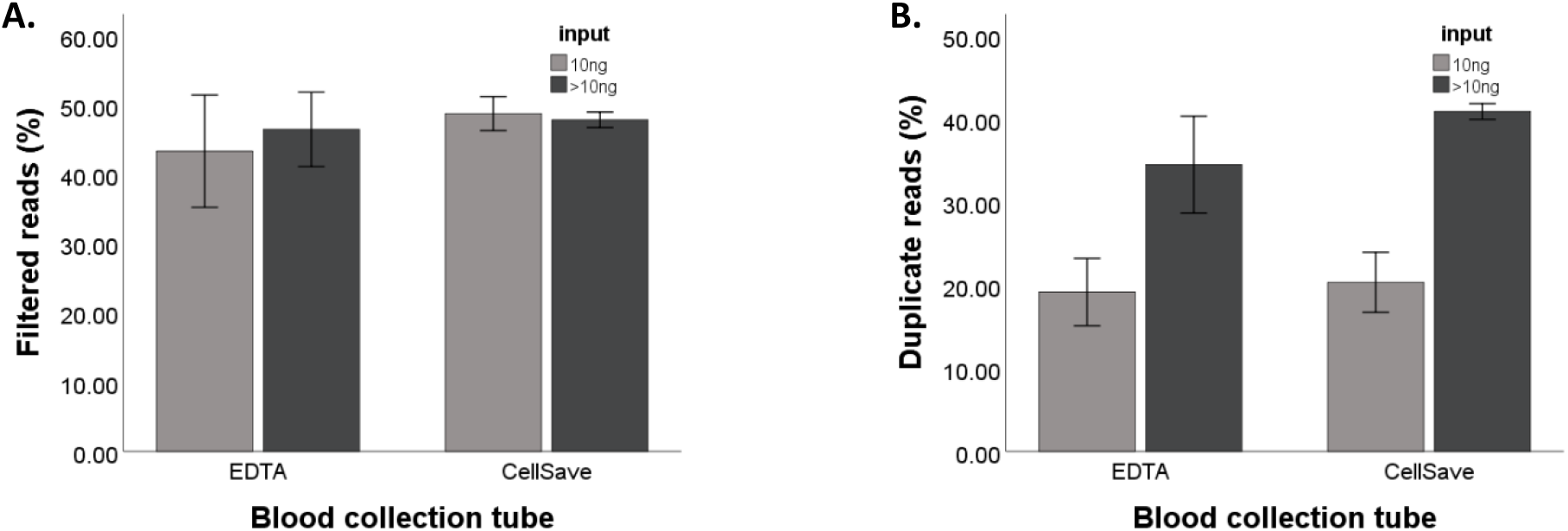
The compatibility of different blood collection tubes with MeD-seq analysis. Percentages of **A**. LpnPI filtered reads and **B**. duplicate reads are shown for EDTA and CellSave tubes obtained during the same blood draw from 3 patients. For each sample 10 ng cfDNA and maximal cfDNA input in 8 µl (>10ng) was analysed, in which the maximal amount was kept equal between EDTA and CellSave per patient.

**Supplementary Figure S2.**
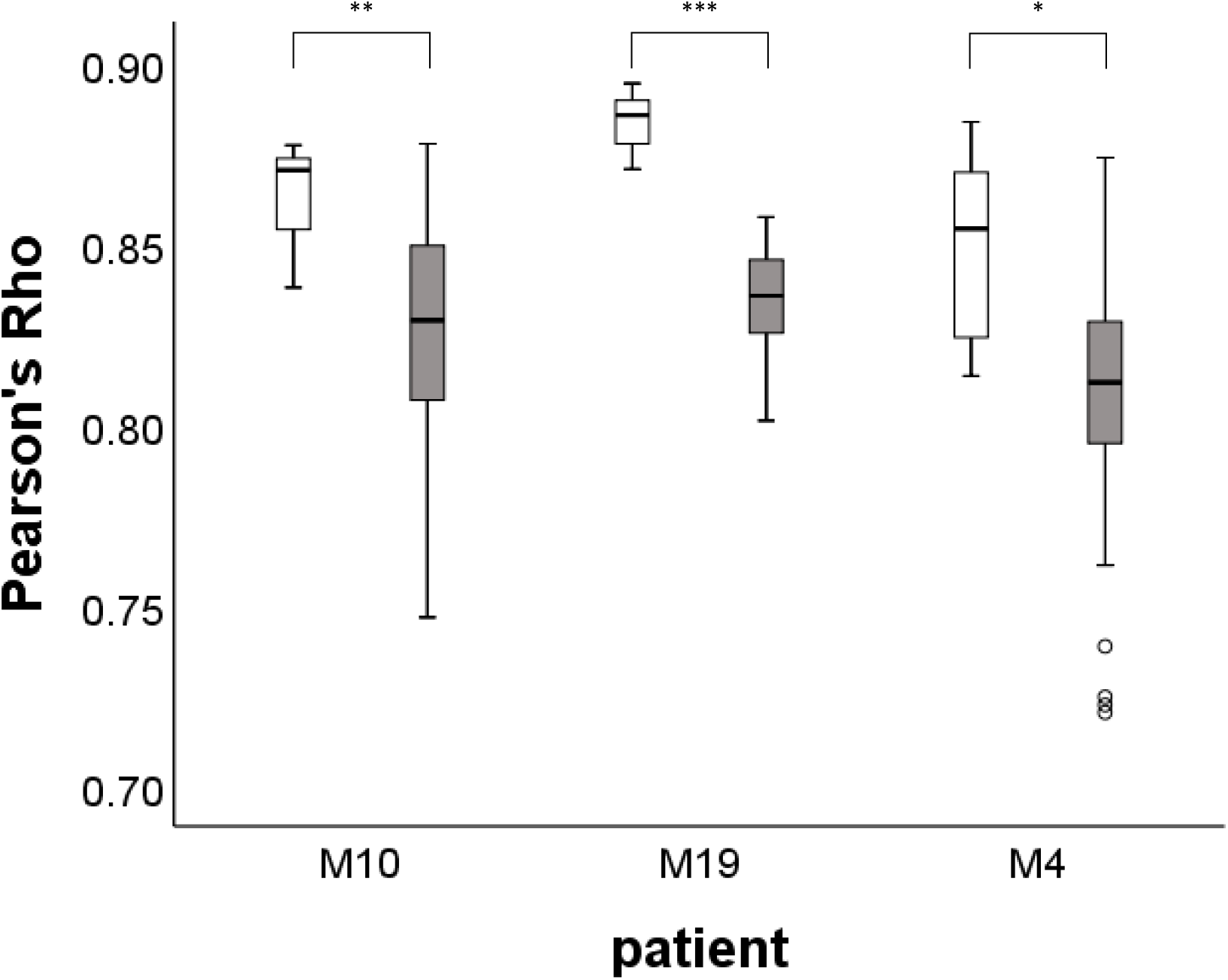
Observed correlations between biological replicates compared to HBDs. Boxplots show significant higher Pearson’s *r* between biological replicates per patient (n=4), in white, compared to the Pearson’s *r* between these samples and unrelated HBD samples (n=9), in gray (Mann-Whitney U, p = 0.015 for M4, p=0.004 for M10, and p<0.001 for M19). Biological duplicates were taken during the same blood draw using either EDTA or CellSave tubes, of which either 10 ng or the maximum yield in 8 µl was used for MeD-seq.

**Supplementary Figure S3.**
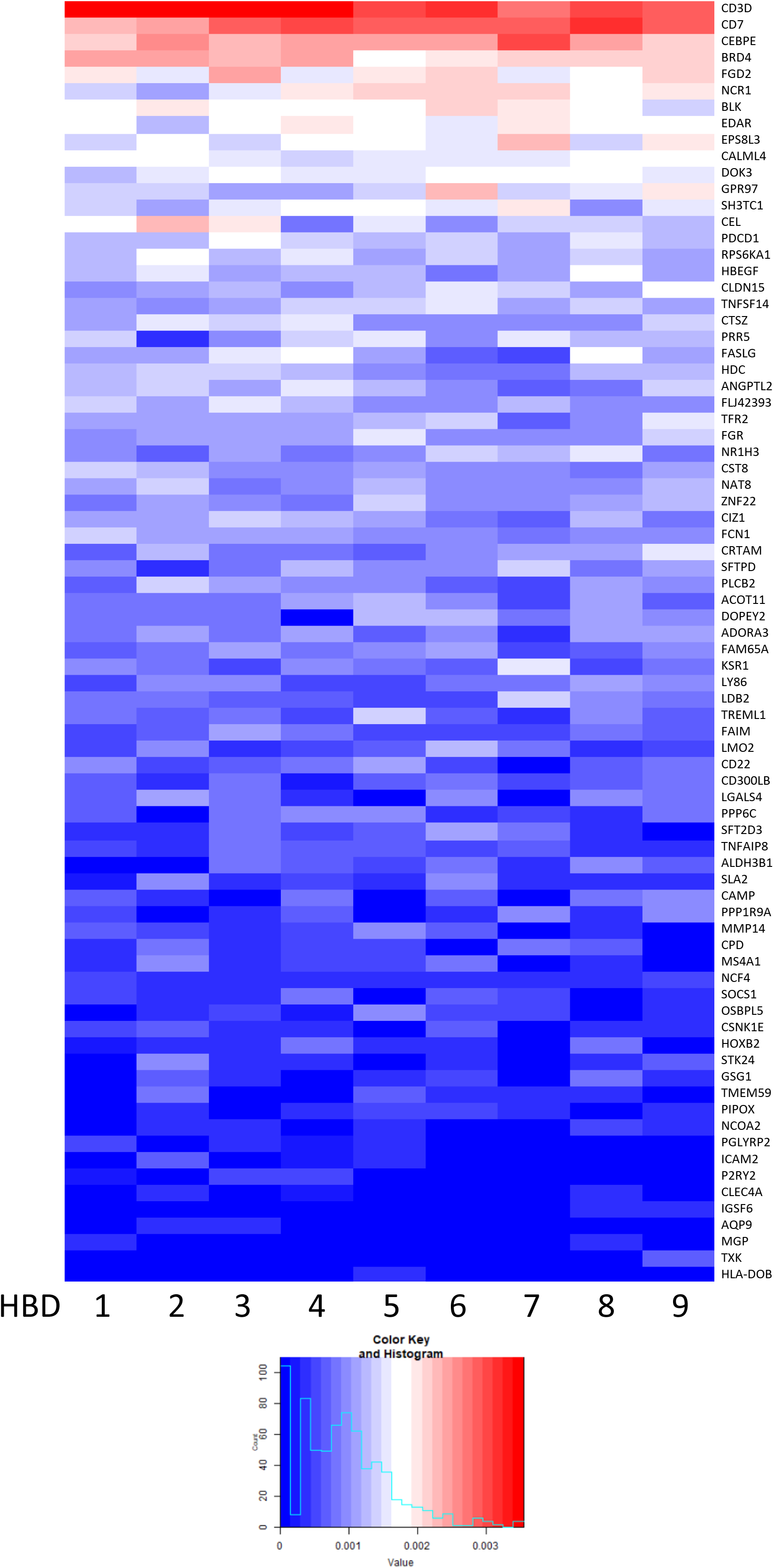
Leukocyte specific markers in HBDs. **A** heat map of a subset of leukocyte specific methylation markers described by Accomando et al.(17) show consistent methylation patterns across the included healthy blood donors. Only regions showing at least 1 read in 9 HBD samples are included.

**Supplementary Figure S4.**
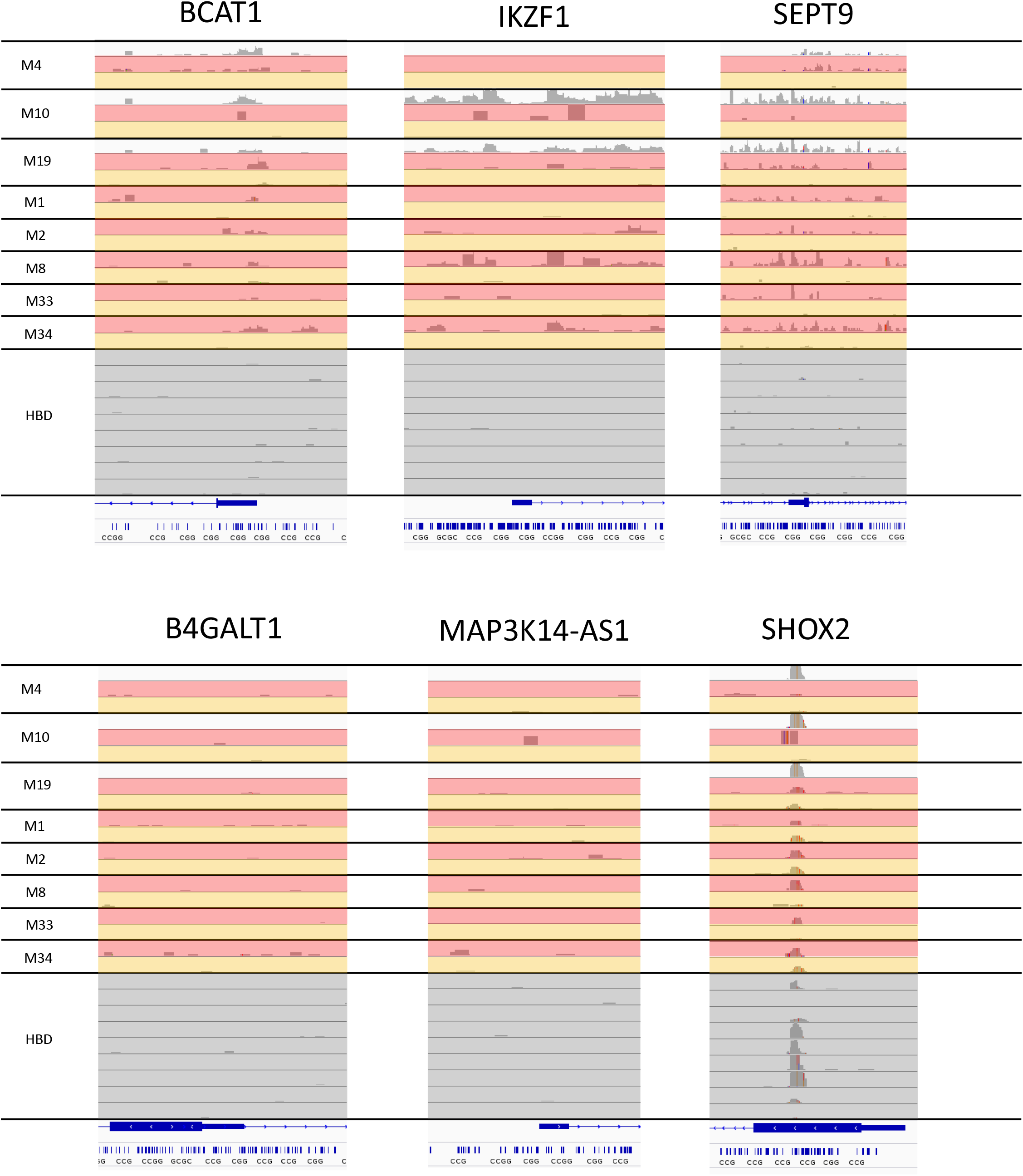
MeD-seq results for published biomarkers on cfDNA from HBDs and paired tissue, pre-operative and post-operative cfDNA from CRLM patients. Sequenced MeD-seq reads shown for BCAT1, IKZF1, SEPT9, B4GALT1, MAP3K14-AS1 and SHOX2 from pre-operative cfDNA (red) and post-operative cfDNA (orange) of CRLM-patients and cfDNA samples of HBDs (black). Results are visualized in Integrative Genomics Viewer (IGV) v.2.9.4.

**Supplementary Figure S5.**
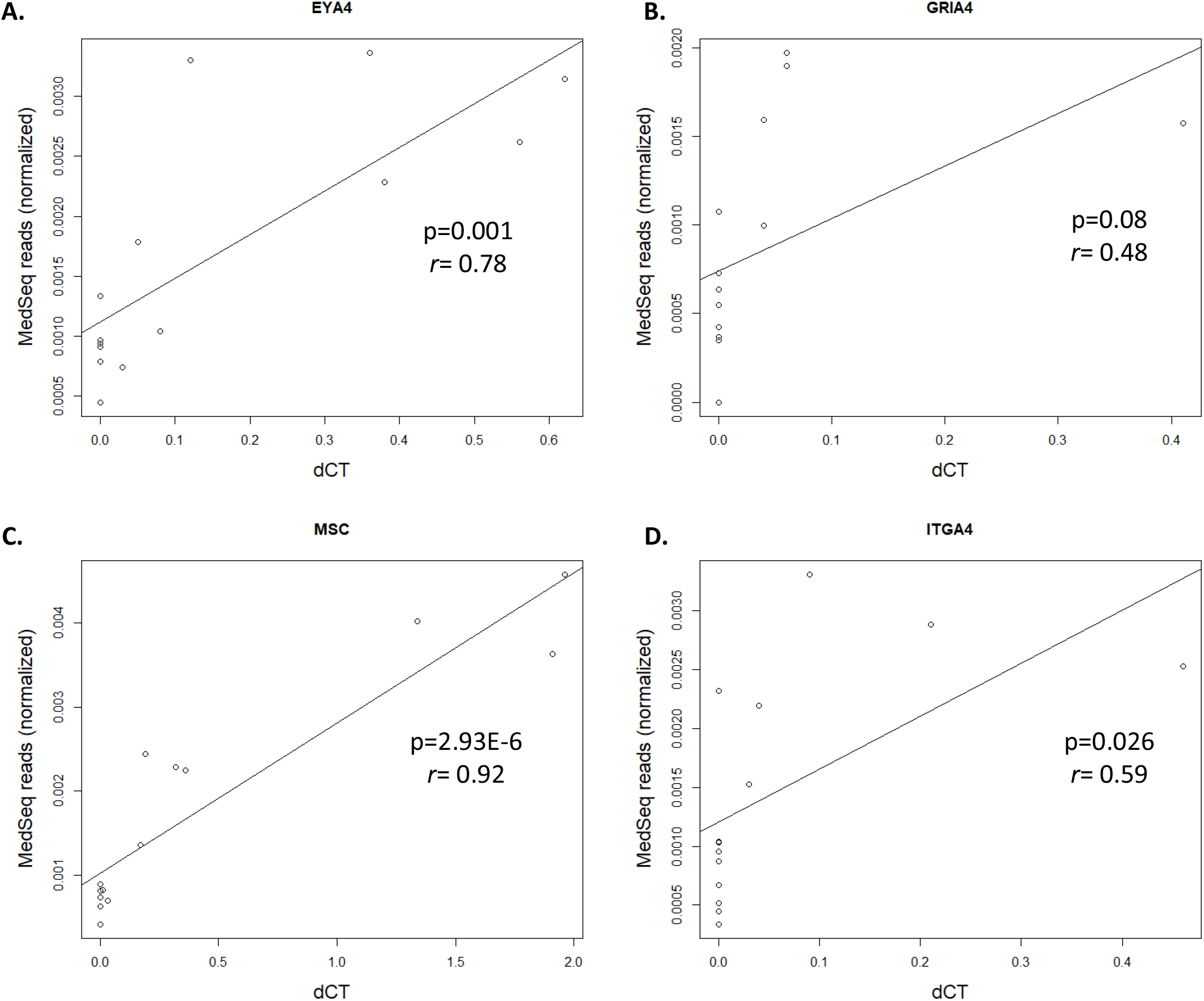
Correlation between MeD-seq and qMSP for previously identified markers for CRLM. Scatterplots showing the Pearson correlations between MeD-seq and qMSP results for **A**. EYA4, **B**. GRIA4, **C**. MSC, and **D**. ITGA4.

